# ECM-derived biophysical cues mediate interstitial flow-induced sprouting angiogenesis

**DOI:** 10.1101/2022.06.04.494804

**Authors:** Chia-Wen Chang, Hsiu-Chen Shih, Marcos Cortes-Medina, Peter E. Beshay, Alex Avendano, Alex J. Seibel, Wei-Hao Liao, Yi-Chung Tung, Jonathan W. Song

## Abstract

Sprouting angiogenesis is orchestrated by an intricate balance of biochemical and mechanical cues in the local microenvironment. Interstitial flow has been established as a potent regulator of angiogenesis. Similarly, extracellular matrix (ECM) physical properties, such as stiffness and microarchitecture, have also emerged as important mediators of angiogenesis. Yet, the interplay between interstitial flow and ECM physical properties in the initiation and control of angiogenesis is poorly understood. Using a 3-D microfluidic tissue analogue of angiogenic sprouting with defined interstitial flow, we found that the addition of hyaluronan (HA) to collagen-based matrices significantly enhances sprouting induced by interstitial flow compared to responses in collagen-only hydrogels. We confirmed that both the stiffness and matrix pore size of collagen-only hydrogels were increased by the addition of HA. Interestingly, interstitial flow-potentiated sprouting responses in collagen/HA matrices were not affected when functionally blocking the HA receptor CD44. In contrast, enzymatic depletion of HA in collagen/HA matrices with hyaluronidase (HAdase) resulted in decreased stiffness, pore size, and interstitial flow-mediated sprouting to the levels observed in collagen-only matrices. Taken together, these results suggest that HA enhances interstitial flow-mediated angiogenic sprouting through its alterations to collagen ECM stiffness and pore size.

## 1. Introduction

Angiogenesis, or the extension or expansion of pre-existing blood vessel networks, is essential for several physiological and pathological processes such as wound healing, inflammation, and cancer^[1]^. During angiogenesis, endothelial cells (ECs) residing in a blood vessel wall undergo morphogenesis to form an angiogenic sprout^[2]^. Newly formed blood vessels by angiogenesis provide nutrients and oxygen to growing tissues^[3]^. Pathologies such as cancer are characterized by uncontrolled angiogenesis that is initiated and supported by cues arising from the host tissue or microenvironment^[4]^. Several microenvironmental cues are known to regulate angiogenesis, including biomolecular signaling, biophysical properties of extracellular matrix (ECM), and biomechanical fluid forces^[5]^. Therefore, because of the broad implications of angiogenesis in health and disease, there is significant interest in understanding the interplay of microenvironmental determinants of EC behavior during neo-vessel outgrowth.

Numerous biomolecules, such as vascular endothelial growth factor (VEGF)^[2a]^, C-X-C motif chemokine 12 (CXCL12)^[6]^, and sphingosine-1-phosphate (S1P)^[7]^, have firmly been established as key controllers of vascular morphogenesis. More recently, fluid mechanical forces associated with blood flow have emerged as important regulators of angiogenesis and vascular functions, either as standalone mediators or as co-determinants of biomolecular-induced sprouting^[8]^. Of these fluid forces, slowly moving fluid flow across the vessel wall due to interstitial plasma flow has been shown to be a potent regulator of angiogenic sprouting^[9]^. Interstitial flow is often elevated in tumor and inflammatory microenvironments due to heightened pressure gradients within the interstitium^[10]^. Interestingly, multiple studies have confirmed that interstitial flow-mediated angiogenic sprouting preferentially occurs when flow is oriented against the direction of sprouting^[11]^ with this process mediated by delocalization of VE-cadherin cell-cell adhesions^[12]^, the small GTPase RhoA^[13]^, αvβ3integrins^[14]^, and other mechanisms.

The composition of the ECM has a profound influence on the biophysical properties of tissue such as mechanical stiffness, microarchitecture, and transport efficiency of nutrients and drug molecules^[15]^. Moreover, dysregulated composition and architecture of the ECM is identified as characteristic of tumors^[16]^, fibrosis^[17]^, and other pathologies. For example, increased levels of fibrillar collagen (e.g., type I and type III) and non-fibrillar components such as proteoglycans and glycosaminoglycans (GAGs) (e.g., hyaluronan or HA) in the tumor ECM are characteristic of aggressive or desmoplastic tumors^[18]^. Recent studies have implicated the composition and physical properties (e.g., stiffness and microarchitecture) of the ECM as important mediators of angiogenesis and microvessel function. For instance, increased tissue stiffness due to matrix cross-linking promotes vessel outgrowth and decreased vessel barrier function^[19]^. Importantly, these responses that were indicative of a tumor vasculature phenotype were independent of VEGF stimulation. Another study determined that the porosity of natural and synthetic hydrogel matrices is a key determinant of angiogenic sprout invasion speed and diameter^[20]^. Recently, our group reported that angiogenesis and vessel permeability responses induced by CXCL12 isoforms were contingent on the type I collagen and HA compositions of the ECM^[21]^.

Despite the wealth of evidence that interstitial flow, ECM composition, and ECM physical properties mediate angiogenesis, to our knowledge, no study has simultaneously examined the interplay of these factors in coordinating endothelial sprouting. Here, we used a biomimetic 3-D microvessel analogue system to systematically measure interstitial flow-potentiated angiogenesis in the context of ECM properties determined by the collagen and HA composition. We observed that the addition of HA to collagen matrices significantly enhanced endothelial sprouting against the direction of interstitial flow. However, under static conditions, addition of HA to collagen-based matrices had no effect on sprouting. Interestingly, interstitial flow-potentiated sprouting responses in collagen/HA matrices were not affected when functionally blocking the HA receptor CD44 with an anti-CD44 antibody. In contrast, enzymatic depletion of HA with hyaluronidase (HAdase) decreased interstitial flow-mediated sprouting in collagen/HA matrices. We confirmed that the stiffness and pore size of collagen-based matrices were increased by the addition of HA. Moreover, in the presence of a broad-spectrum matrix metalloproteinase (MMP) inhibitor (GM6001), interstitial flow-mediated sprouting was significantly higher in collagen/HA versus collagen-only matrices. These results suggest that HA enhances interstitial flow-mediated angiogenic sprouting through its alterations to ECM stiffness and pore size. Collectively, our study provides novel insights into the role of the biophysical properties of the ECM in controlling angiogenesis in the presence of interstitial flow.

## 2. Results

### 2.1 HA enhances interstitial flow-mediated sprouting angiogenesis in collagen-based matrices

A biomimetic microfluidic microvessel analogue system was fabricated using poly(dimethylsiloxane) (PDMS) soft lithography to study the combined effects of interstitial flow and ECM composition on sprouting angiogenesis (**Figure 1A**). The configuration of the microfluidic device was composed of three parallel microchannels (**Figure 1B**). The two outer channels were denoted as “vascular channel” and “fluidic channel”. The vascular and fluidic channels flank a localized ECM region (See “Materials and Methods” for the dimensions of the engineered microvessel analogue). The vascular channel was fully lined with human umbilical vein endothelial cells (HUVECs) to comprise a microvessel analogue (**Figure 1C**). The fluidic channel provided fluid pressure source for the generation of interstitial flow of ∼11 µm/s (**Figure 1D**) (See Materials and Methods). Consequently, interstitial flow was oriented from the fluidic channel, through the ECM region, and into the vascular channel. With this orientation, interstitial flow was applied against endothelial sprouting from the vascular channel. This interstitial flow orientation was used because numerous studies have shown that endothelial cells preferentially sprout against or opposite the direction of interstitial flow^[22]^. Interstitial flow is estimated to be at a level of ∼1 μm/s in normal tissue^[23]^. This flow velocity is substantially increased in pathological conditions such as inflammation and cancer^[5d, 24]^. The levels of flow that we impose are in line with previous reports in 3-D culture models that reflect pathological values of interstitial flow velocity^[14, 25]^. The central channel was filled with either Type I collagen ECM (henceforth referred to as “collagen-only”) or a mixture of type I collagen gel and HA^[26]^ (henceforth referred to “collagen/HA”). The concentration of 3 mg/mL Type I collagen was used for both collagen-only and collagen/HA ECM in this study. For collagen/HA conditions, HA at a concentration of 1 mg/mL was mixed with collagen gel prior to polymerization. The ratio of collagen to HA used in this study is relevant to tumor physiology as it is in line with the collagen/HA composition of breast tumors *in vivo*^[27]^.

**Figure 1.**
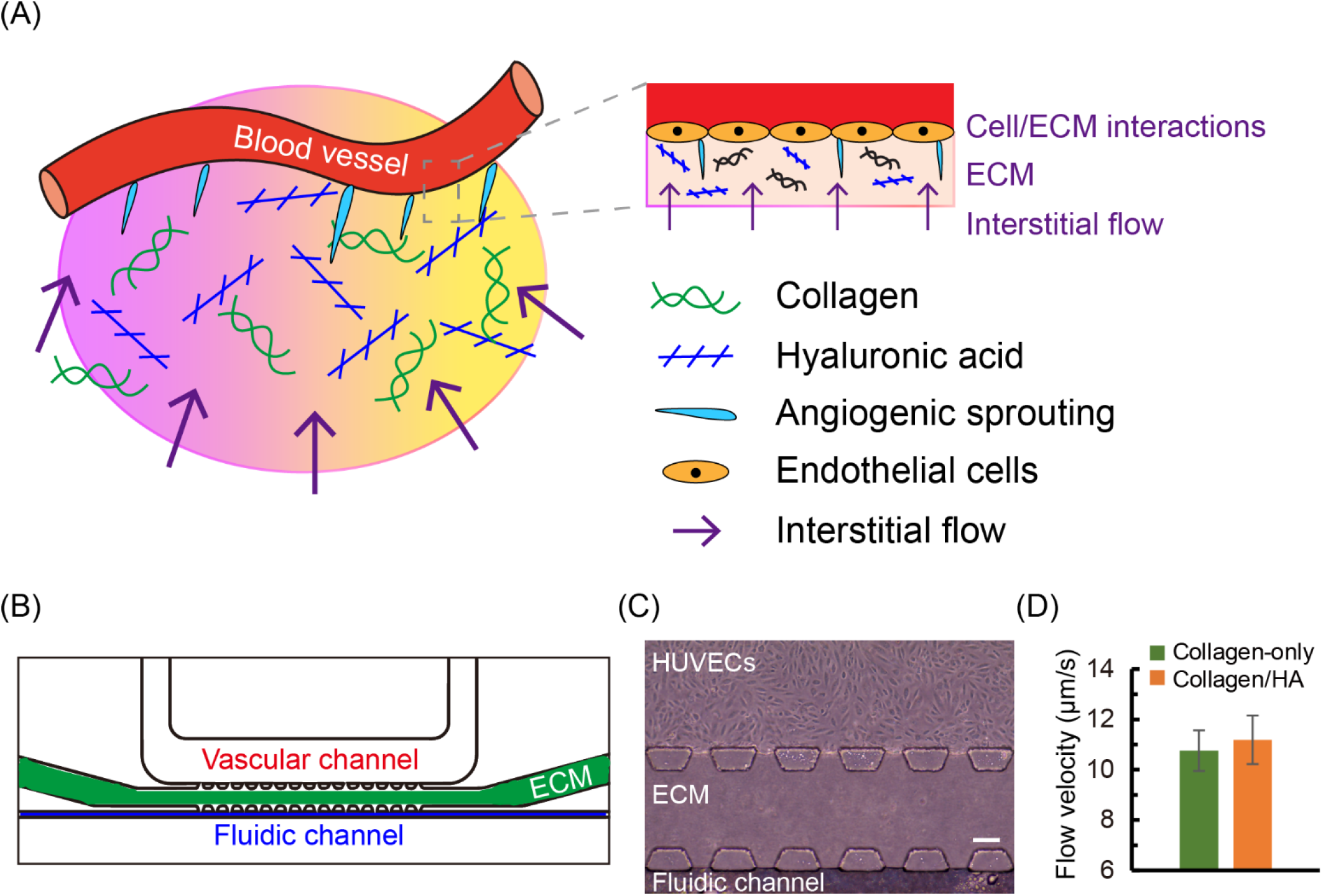
3-D biomimetic microfluidic microvessel model for studying interstitial flow-mediated endothelial cells sprouting. (A) Schematic of interstitial flow-mediated angiogenic sprouting in tissue microenvironment; (B) Schematic of developed microfluidic microvessel analogue for studying interstitial flow-induced angiogenic sprouting; (C) Phase-contrast image of human umbilical vein endothelial cells (HUVECs) cultured in microfluidic microvessel analogue. Scale bar = 80 μm; (D) Interstitial flow velocity in microvessel of collagen-only and collagen/HA ECM. The data were expressed as mean ± standard error of mean (n=5 for collagen-only gel; n=7 for collagen/HA matrices).

To identify the effects of ECM composition on angiogenic sprouting, we first compared sprouting in collagen-only ECM and collagen/HA matrices under static conditions using our HUVEC-lined microfluidic vessel analogue system. HUVECs did not show observable sprouting in both collagen-only ECM and collagen/HA matrices after three days of static culture (**Figure 2 and Figure S1**). These results demonstrate that introduction of HA into the collagen-based ECM did not prompt HUVEC sprouting under static conditions. Typically, angiogenesis is coincident with increased vessel permeability,^[4]^ and we previously showed that introduction of HA into collagen-based ECM stabilizes vessel barrier function under static conditions. In contrast, interstitial flow applied against the endothelium of the vessel channel induced HUVEC sprouting for both the collagen-only and the collagen/HA matrices (**Figure 3**). For collagen-only ECM, the average sprouting area increased from 914 µm^2^ on day 1 to 3429 µm^2^ on day 3. For collagen/HA ECM, the average sprouting area increased from 1835 µm^2^ on day 1 to 5158 µm^2^ on day 3 (**Figure 3A**). These results confirm that interstitial flow promotes angiogenesis when applied against the direction of sprouting. However, it is notable that interstitial flow significantly increased HUVEC sprouting for the collagen/HA matrices by ∼2-fold compared to the collagen-only ECM on day 1. These results suggest that addition of HA to collagen-based ECM promotes sprouting of the microvessels in the presence of interstitial flow (**Figure 3A**).

**Figure 2.**
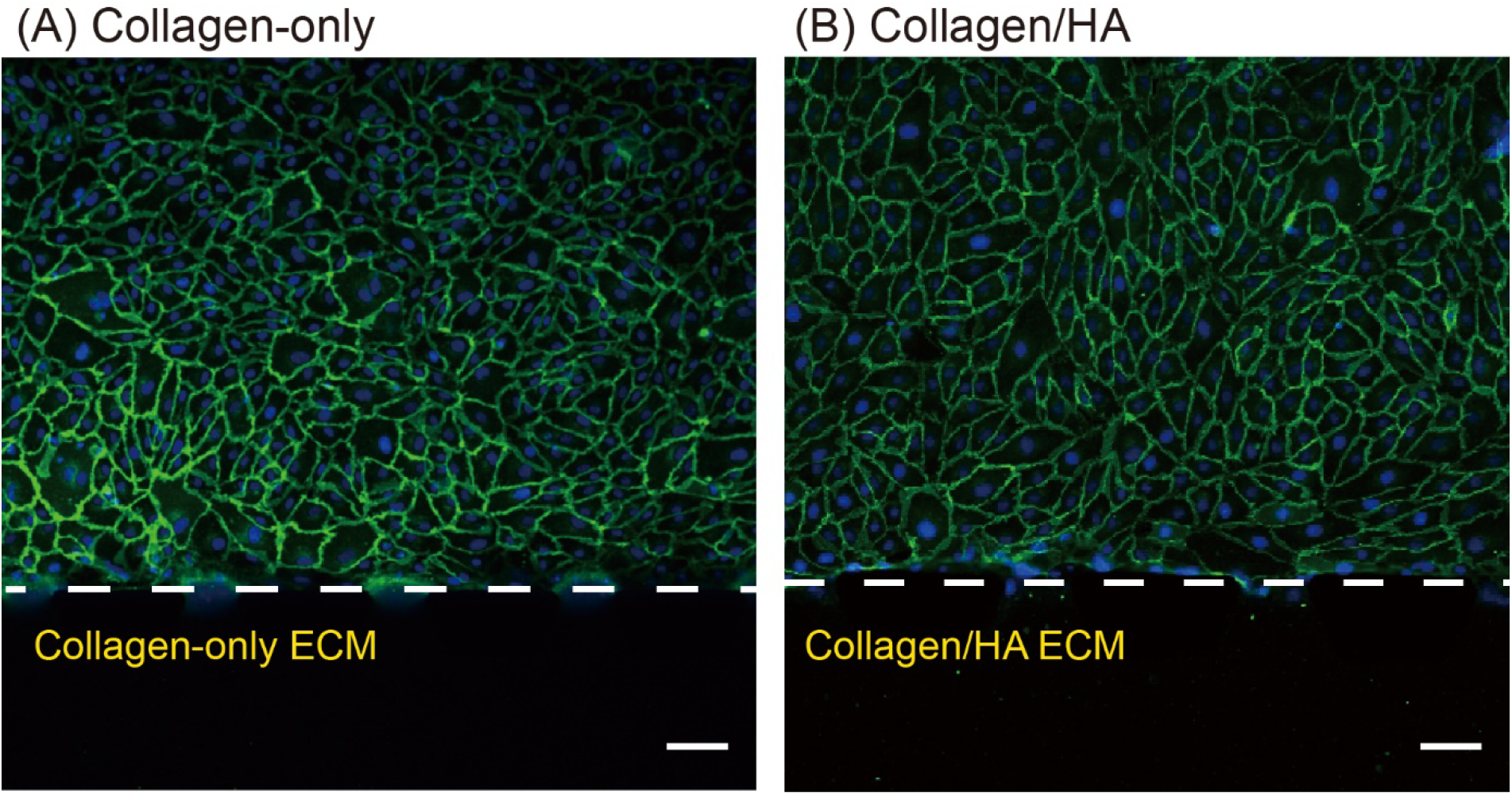
Endothelium of HUVECs in the microfluidic microvessel analogue under static condition on day 3. VE-cadherin junction protein (green) and DAPI (blue) staining of microvessels in (A) collagen-only and (B) collagen/HA matrices under static condition. Dash white lines indicate the ECM/Cell interfaces. (Scale bars = 50 µm)

**Figure 3.**
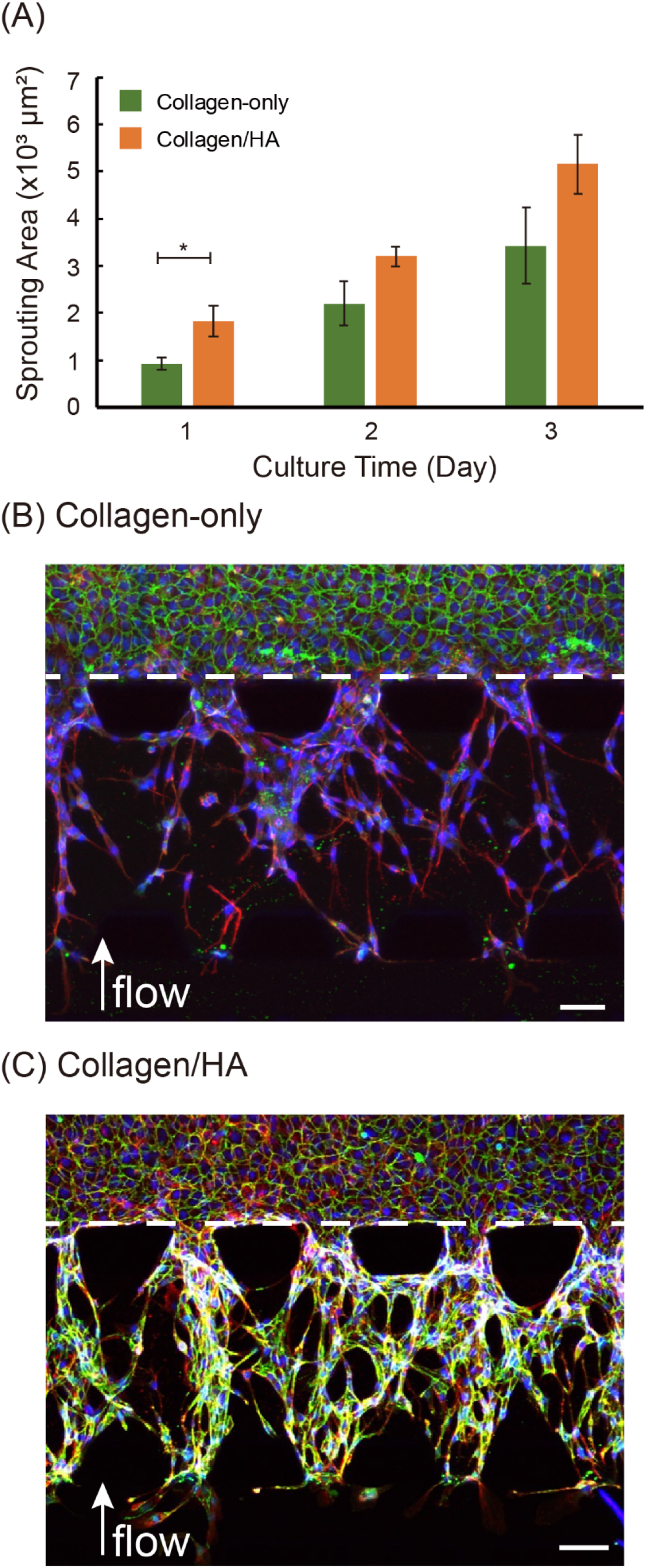
Sprouting angiogenesis mediated by interstitial flow in collagen-only and collagen/HA matrices. (A) sprouting area of microvessels in both collagen-only and collagen/HA matrices; The morphology of sprouting in the microvessel analogue is assessed by staining VE-cadherin junction protein (green), F-actin (red) and DAPI (blue) of (B) collagen-only and (C) collagen/HA metrices. White arrow indicates the direction of interstitial flow. The data were expressed as mean ± standard error of mean (n=5). One-way ANOVA followed by post-hoc unpaired, two-tailed Student t test was performed to evaluate the statistical significance. * indicates p-value < 0.05. Dash white lines indicate the ECM/Cell interfaces. (Scale bar = 50 µm)

### 2.2 HA promotes interstitial flow-induced sprout elongation in collagen-based matrices

Interstitial flow has been shown to be necessary for sustaining endothelial sprout elongation in collagen-only ECM^[8b]^. Interestingly, we observed increased sprout elongation in response to interstitial flow in collagen/HA ECM versus collagen-only ECM. Namely, at day 3, some of the HUVEC sprouts in the collagen/HA ECM reached the distal fluidic channel while HUVEC sprouts in the collagen-only ECM remained within the ECM compartment (**Figure 3B and 3C**). These observations prompted us to compare interstitial flow-mediated sprout elongation in the collagen-only and collagen/HA matrices over multiple days. For this comparison, we set 10 equally divided regions of the ECM. Each section was assigned a number ranging from 1 (proximal endothelium/ECM interface) to 10 (distal fluidic channel) (**Figure 4A**). On day 1, interstitial flow-promoted sprouting in the collagen/HA ECM was more advanced compared to the collagen-only ECM. For the collagen/HA ECM, most of the sprouting was in sections 2 - 4 while most of the sprouting for the collagen-only ECM was distributed in sections 1 - 2 (**Figure 4B**). Sprout elongation in the collagen/HA ECM continued to be more advanced compared to the collagen-only ECM for days 2 and 3 (**Figure 4C and 4D**). These results imply that the addition of HA to collagen-based ECM enhances endothelial sprout elongation in response to interstitial flow over the observed timeframe of 3 days.

**Figure 4.**
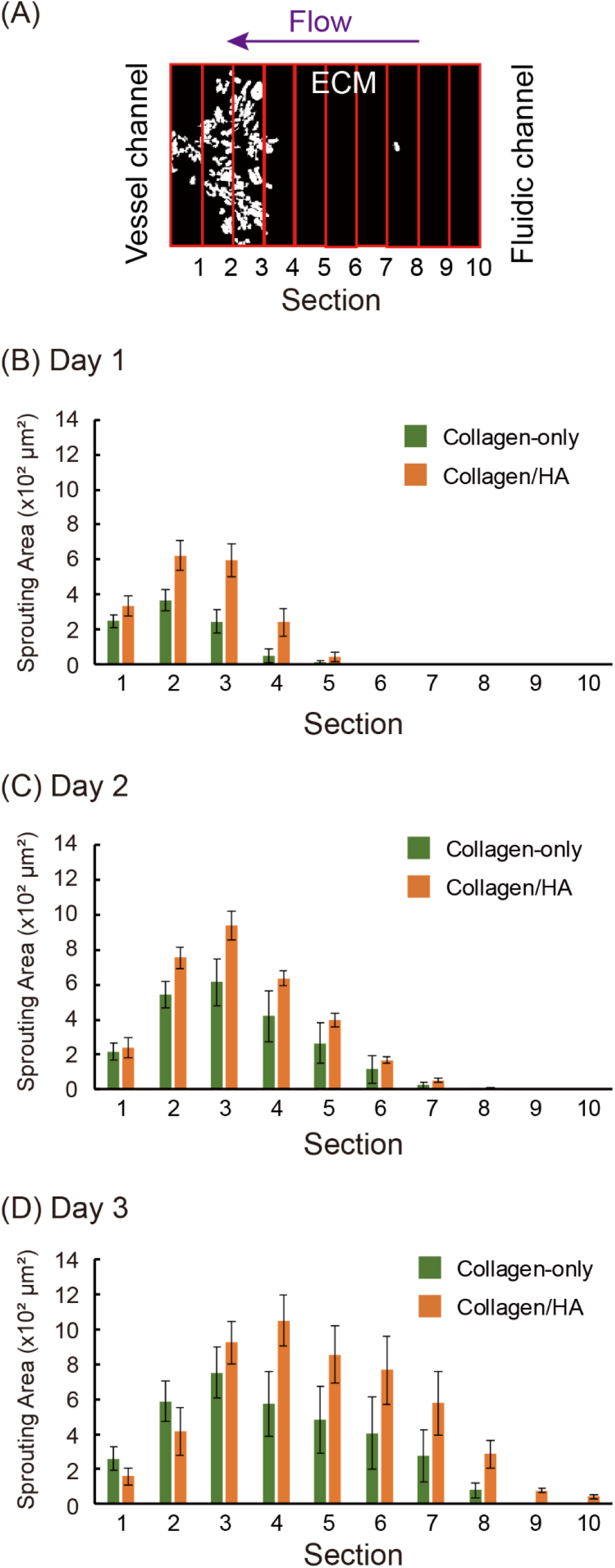
Collagen/HA matrices promote the elongation of angiogenic sprouting in interstitial-flow mediated sprouting angiogenesis. (A) The schematic of each section in ECM compartment. The distribution of angiogenic sprouting in both collagen-only and collagen/HA matrices of entire ECM compartment on (B) Day 1, Day 2 and (D) Day 3. The data were expressed as mean ± standard error of mean (n=5).

### 2.3 Interstitial flow-induced sprouting is partially dependent on MMP activity

Furthermore, we assessed the role of matrix metalloproteinases (MMPs) on interstitial flow-induced sprouting into collagen-only and collagen/HA matrices. MMPs are known regulators of sprouting angiogenesis by cleaving peptide bonds of ECM proteins, including collagens^[28]^. In addition, MMPs have been shown to be involved in interstitial flow-mediated HUVEC sprouting in collagen-only ECM.^[8a, 29]^ We tested the effect of inhibiting MMPs with a full spectrum inhibitor of MMP activity (20 μM, GM6001, Millipore Sigma). Inhibition of MMP activity could partially impede interstitial flow-mediated sprouting in both collagen-only and collagen/HA matrices (**Figure 5A and 5B**). However, in the presence of GM6001, the normalized number of sprouts percentage in the collagen/HA ECM (105.2 %) was significantly higher than that in the collagen-only ECM (82.5%). (See Materials and Methods for the normalized number of sprouts percentage) (**Figure 5C)**. Similarly, in the presence of GM6001, the average sprouting area from the microvessel analogue into the collagen/HA ECM (765 µm^2^) was also significantly higher than the sprouting area into the collagen-only ECM (467 μm^2^) on day 1. (**Figure 6**). In addition, the sprouting area for the GM6001 treated microvessels was significantly less for both the collagen-only and collagen/HA matrices compared to their respective interstitial flow controls (**Figure 6**). Our results for the GM6001 treated microvessels agree with the observation from a previous report that interstitial flow-induced sprouting is partially dependent on MMP activity^[8a]^.

**Figure 5.**
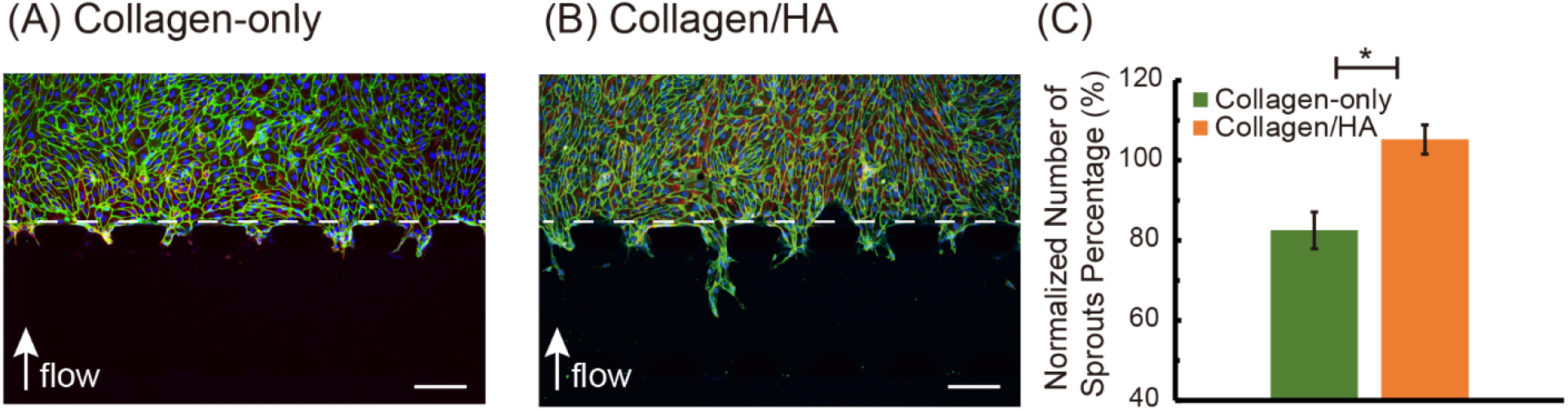
VE-cadherin junction protein (green), actin (red) and DAPI (blue) staining of GM6001-treated angiogenic sprouting of the microvessel analogues in (A) collagen-only; (B) collagen/HA matrices on day 1 culture; (C) Normalized number of sprouts precentage of GM6001-treated HUVECs sprouts of microvessels in both collagen-only and collagen/HA matrices. White arrow indicates the direction of interstitial flow. The data were expressed as mean ± standard error of mean (n=3). One-way ANOVA followed by post-hoc unpaired, two-tailed Student t test was performed to evaluate the statistical significance. * indicates p-value < 0.05. (Scale bars = 150 µm)

**Figure 6.**
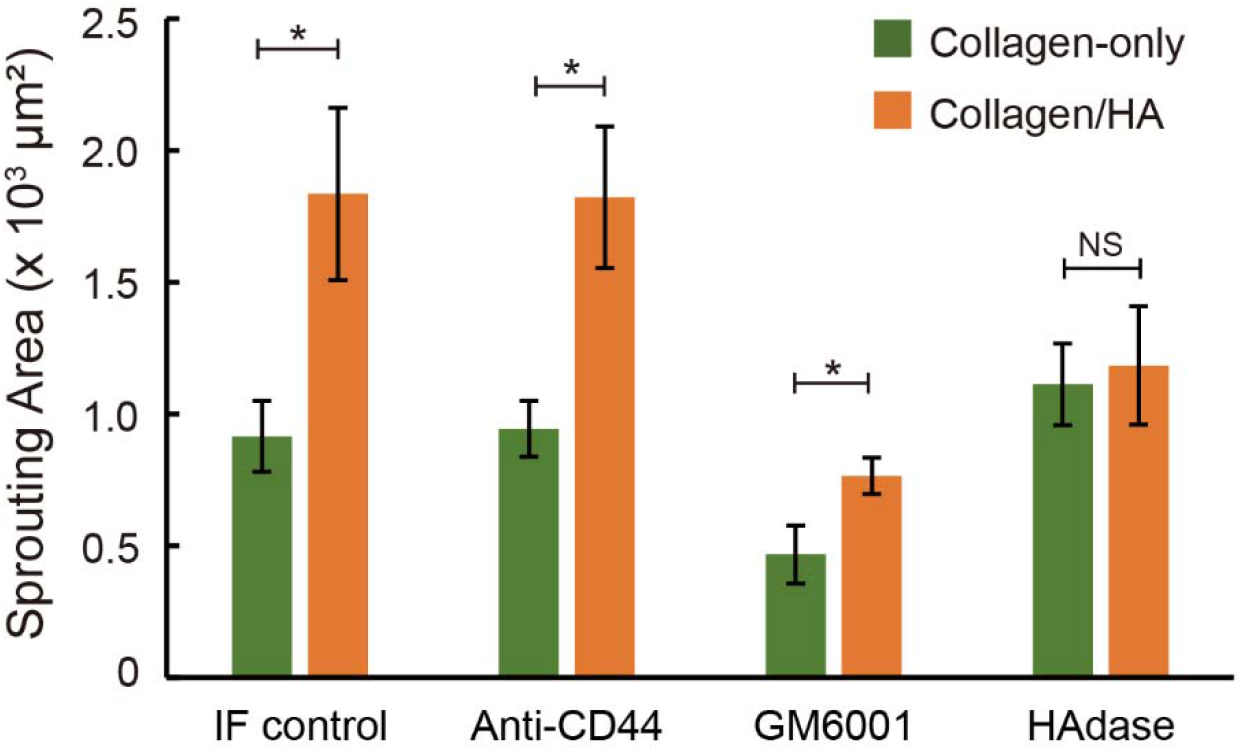
Sprouting area of HUVECs cultured in microvessel analogues of both collagen-only and collagen/HA matrices in the responses of various pharmaceutical drugs treatments (i.e. CD44 blocking, MMP inhibition and enzymatic matrix degradation) under interstitial flow on day 1. The data were expressed as mean ± standard error (n ≥ 3 for all experimental conditions). One-way ANOVA followed by post-hoc unpaired, two-tailed Student t test was performed to evaluate the statistical significance. * indicates p-value < 0.05 and NS indicates no significant difference.

### 2.4 Hyaluronidase but not anti-CD44 inhibits interstitial flow-induced sprouting in collagen/HA matrices

The observed enhancement in angiogenic sprouting due to HA and in the context of interstitial flow led us to investigate the involvement of HA-mediated signaling and biophysical properties. To examine the role HA-mediated signaling, we used a blocking antibody for the HA receptor CD44 (2 μg/mL, Invitrogen, MA5-13890). Endothelial CD44 has been shown to be involved with angiogenesis *in vivo*^[30]^ although its role in interstitial flow-promoted sprouting is not known. For collagen-only ECM and in the presence of interstitial flow, anti-CD44 treatment did not significantly affect sprouting activity (943 µm^2^ versus 914 µm^2^ for anti-CD44 and untreated control respectively) (**Figure 6**). Surprisingly, in collagen/HA ECM, anti-CD44 treatment in the presence of interstitial flow did not affect vessel sprouting (1822 µm^2^ versus 1835 µm^2^ for anti-CD44 and untreated control respectively). These results suggest that endothelial CD44 signaling was not essentially involved in interstitial flow-potentiated sprouting in collagen/HA matrices. We also tested the effect of co-application of MMP inhibitor (GM6001) and anti-CD44 (**Figure S2**, Supplemental Information). Compared to GM6001-only treated microvessels, co-application of GM6001 with anti-CD44 did not significantly reduce sprouting in the presence of interstitial flow for both the collagen-only and collagen/HA matrices. These results suggest that sprouting inhibition in the presence of interstitial flow due to the combinatorial treatment of GM6001 and anti-CD44 was attributed to inhibition of MMP activity and not disrupting engagement of endothelial CD44 with ECM HA.

Moreover, we examined whether the physical presence of HA enhances interstitial flow-mediated sprouting. To conduct these studies, we enzymatically degraded HA with 1 mg/mL hyaluronidase (HAdase, Sigma). HAdase, namely its PEGylated human recombinant form (PEGPH20), has been tested in preclinical and clinical settings for treating stroma-rich, desmoplastic tumors such as pancreatic ductal adenocarcinoma (PDAC)^[31]^. In addition, our group has previously shown that HAdase alleviates barriers to convective drug transport through the interstitial matrix by nullifying extracellular HA synthesis by stromal fibroblasts^[32]^. Regarding endothelial mechanobiology, HAdase has been used to selectively deplete HA of the endothelial glycocalyx, which resulted in blocking increases in vessel permeability induced by intravascular shear stress^[33]^. However, the role of HAdase in mediating angiogenesis, especially in the context of interstitial flow, is unclear. Upon treatment with HAdase, the sprouting area significantly decreased (36%) in the collagen/HA ECM and in the presence of interstitial flow compared to the untreated condition (**Figure 6**). These results confirm that the addition of HA significantly enhances interstitial flow-mediated angiogenesis, which can be nullified by HAdase treatment. Notably, the sprouting area for HAdase treated microvessels in the collagen/HA ECM (1183 µm^2^) was comparable to both the HAdase-treated collagen-only ECM (1112 µm^2^) and untreated collagen-only control (914 µm^2^) (**Figure 6**). These results show that inhibition of interstitial flow-mediated sprouting by HAdase is contingent on HA present in the ECM.

### 2.5 HAdase but not GM6001 alters the mechanical and structural properties of collagen/HA matrices

Next, we investigated the effects of GM6001 and HAdase treatment on the biophysical properties of acellular ECM hydrogel mixtures (i.e., collagen-only and collagen/HA). These studies were motivated by our results that GM6001 and HAdase, but not anti-CD44 treatment, significantly inhibits interstitial flow-mediated sprouting of the HUVEC microvessel analogues in collagen/HA ECM (**Figure 6**). We hypothesized that alterations to biophysical properties due to the addition of HA to collagen-based matrices mediate interstitial flow promoted sprouting. The biophysical properties of the ECM that we characterized were the mechanical stiffness and the microstructural parameter of mean matrix pore size. We characterized these ECM properties because they have been implicated in regulating endothelial sprouting, although under static culture conditions only^[19-20]^. To characterize the stiffness profiles of the acellular ECM hydrogel mixtures, we used indentation testing, as previously described by our group.^[21, 34]^ To characterize the mean matrix pore size, we employed confocal reflectance microscopy and image analysis techniques described previously by our group^[34]^.

We first benchmarked changes in mechanical and microstructural properties of untreated collagen-only and collagen/HA matrices. The measured stiffness for the 3 mg/mL collagen-only gel was 3.15 kPa (**Figure 7**). Addition of HA (1 mg/mL) to the collagen-only gel increased the measured stiffness by approximately 2-fold to 6.44 kPa (**Figure 7**). These stiffness measurements were in line with our previous reports for the same ECM compositions.^[21, 34]^ Moreover, addition of HA to the collagen-based ECM significantly increased the mean pore size by 13% from 1.70 µm to 1.92 µm compared to collagen-only ECM (**Figure 8**). We previously reported a similar outcome on matrix pore size when comparing the same ECM compositions^[34]^. When integrating the endothelial sprouting results (**Figure 6**) with the biophysical measurements (**Figures 7 and 8**), our findings show that the enhancement in interstitial flow-promoted sprouting due to addition of HA to collagen-based ECM correlates with HA-induced increases in ECM stiffness and matrix pore size.

**Figure 7.**
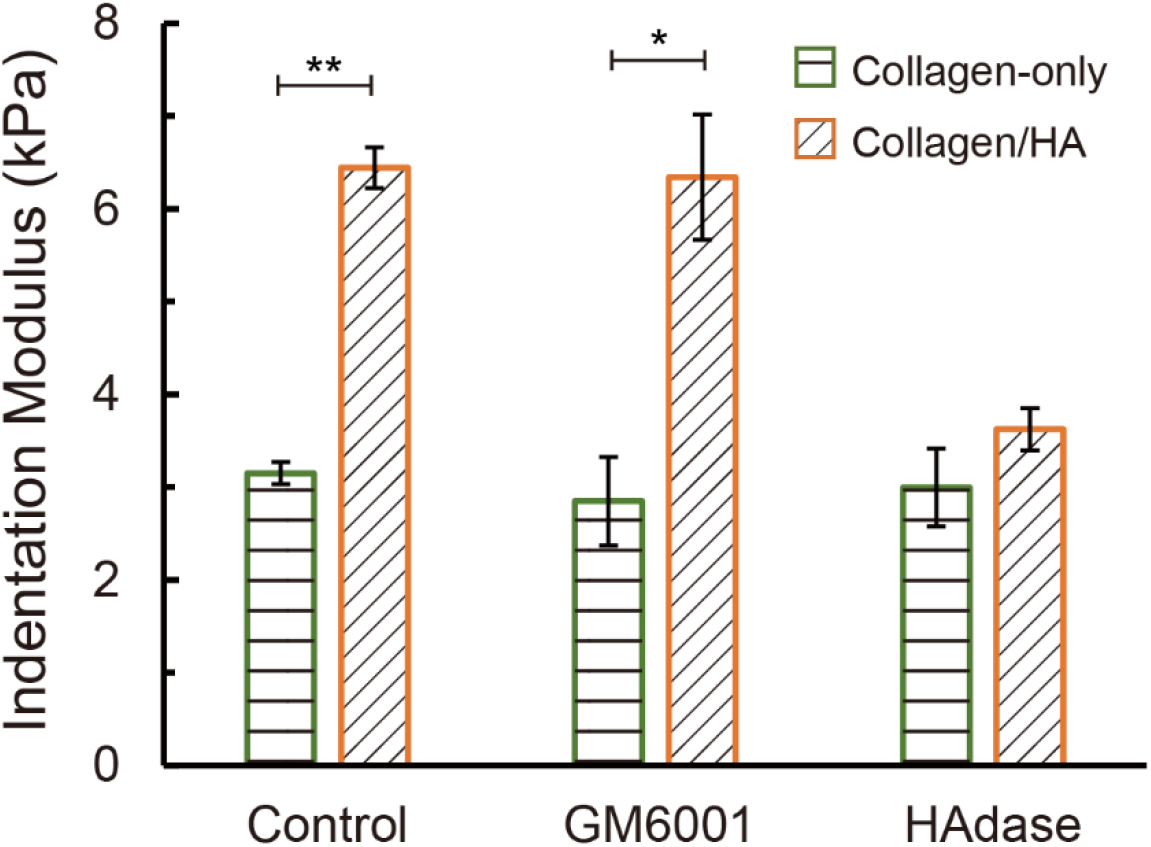
ECM characterizations: indentation modulus of both collagn-only and collagen/HA matrix in various experimental conditions (MMP inhibition and enzymatic matrix degradation). The biochemical treatment was applied for 24 hours before indentation testing was performed. The data were expressed as mean ± standard error of mean (n ≥ 3 for all experimental conditions). One-way ANOVA followed by post-hoc unpaired, two-tailed Student t test was performed to evaluate the statistical significance. * and ** indicate p-value < 0.05 and < 0.01, respectively.

**Figure 8.**
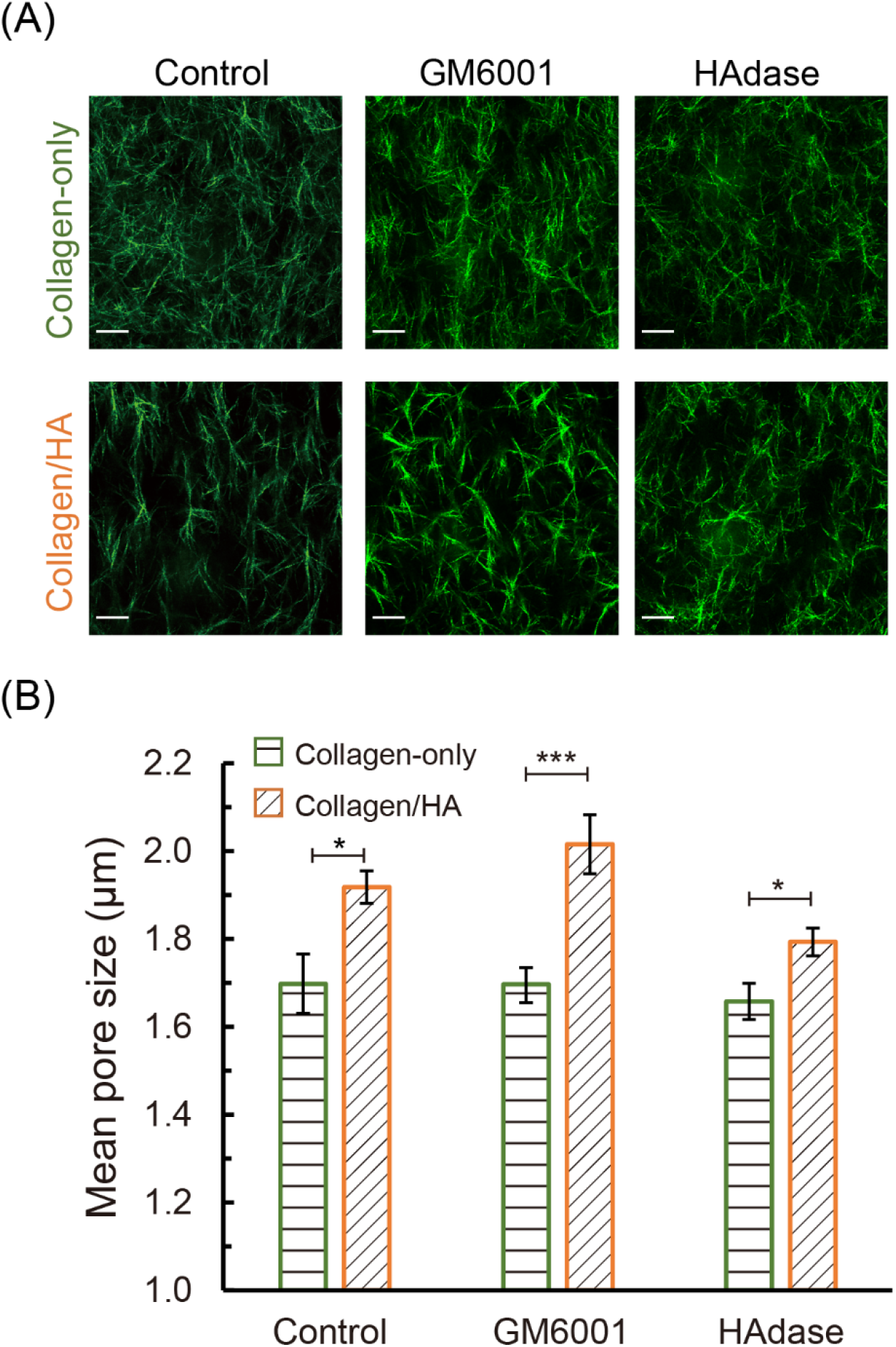
ECM characterizations: (A) Reflectance confocal microscopic images of both collagen-only and collagen/HA matrix in various experimental conditions (MMP inhibition and enzymatic matrix degradation). The biochemical treatment was performed for 24 hours prior to confocal microscopy imaging; (B) The quantification of matrix pore size in various experimental conditions at static (n ≥ 3 for all experimental conditions). The data were expressed as mean ± standard error of mean. One-way ANOVA followed by post-hoc unpaired, two-tailed Student t test was performed to evaluate the statistical significance. *, ** and *** indicate p-value < 0.05, p < 0.01 and p < 0.001, respectively. (Scale bars = 20 µm)

Subsequently, we confirmed that addition of GM6001 to acellular collagen-only and collagen/HA did not significantly change the stiffness measurements for collagen-only (2.85 kPa) and collagen/HA (6.34 kPa) compared to the respective untreated control conditions (3.15 kPa and 6.44 kPa) (**Figure 7**). In addition, GM6001 did not significantly alter the mean matrix pore size for both collagen-only (1.70 µm) and collagen/HA matrices (2.06 μm) compared to the corresponding untreated conditions for these ECM hydrogel mixtures (1.70 µm for collagen-only and 1.92 µm for collagen/HA, respectively) (**Figure 8**). Since GM6001 is an inhibitor of cell secreted MMPs, we did not expect this pharmacological agent to alter the mechanical and structural properties of acellular matrices. These results imply that partial inhibition of interstitial flow-potentiated endothelial sprouting by GM6001 (**Figure 6**) was due to blocking MMP activity and not attributed to direct modifications to the ECM mechanical and structural properties.

Finally, we characterized the mechanical and structural ECM properties due to HAdase treatment. For the collagen-only ECM, as expected, we observed no significant changes in the stiffness profile 3.00 kPa (**Figure 7**) and mean matrix pore size (1.66 μm) (**Figure 8**) due to HAdase treatment compared to untreated collagen-only ECM. In contrast, treatment of the collagen/HA ECM with HAdase significantly decreased both the stiffness by 43% from 6.44 kPa to 3.62 kPa and mean matrix pore size by 7% from 1.92 μm to 1.79 μm (**Figures 7 and 8**). Moreover, the stiffness of HAdase-treated collagen/HA ECM (3.62 kPa) was comparable to HAdase-treated collagen-only ECM (3.00 kPa) and untreated collagen-only ECM (3.15 kPa). Similarly, the mean matrix pore size of HAdase treated collagen/HA ECM (1.79 μm) was comparable to HAdase-treated collagen-only ECM (1.66 μm) and untreated collagen-only ECM (1.70 μm). Therefore, treatment of collagen/HA ECM with HAdase significantly decreased both stiffness and mean matrix pore size to levels observed for the collagen-only ECM (both untreated and HAdase treated). These results suggest that the selective decrease in interstitial flow-promoted sprouting due to HAdase treatment in collagen/HA matrices (**Figure 6**) may be attributed to HAdase decreasing the stiffness and mean matrix pore size of this ECM hydrogel mixture (**Figures 7 and 8**).

## 3. Discussion

Composition of the 3-D ECM profoundly alters biophysical properties of tissue, such as mechanical stiffness, microarchitecture, and transport efficiency of nutrients and drug compounds^[15]^. While biophysical properties of tissue can be measured *in vivo*, studying the effects of these properties on angiogenesis *in vivo* remain challenging due to limited experimental control over ECM composition. To overcome this fundamental challenge to studying ECM, our design criteria included rigorous control and detailed characterization of the biophysical properties of the reconstituted ECM-derived hydrogels introduced in our *in vitro* microtissue analogue system for studying angiogenesis. This approach enabled us to define how physical properties of the ECM regulate angiogenesis that is potentiated by interstitial flow.

Our study emphasizes angiogenesis responses due to modifications of the collagen ECM by HA addition. We demonstrate that the pro-angiogenic consequences of HA addition to collagen ECM modifications are realized in the presence of interstitial flow only. Interestingly, we observe that functionally blocking HA-CD44 interactions with anti-CD44 did not affect interstitial flow-induced sprouting in collagen/HA ECM. This result is in contrast with our previously reported observation that under static conditions, neutralizing HA-CD44 interactions with anti-CD44 completely restored endothelial sprouting levels in collagen/HA matrices compared to the collagen-only counterpart^[21]^. These results suggest that while CD44 is key for controlling HA-mediated angiogenic responses under static conditions, it is not essentially involved in angiogenic responses induced by interstitial flow. This outcome is surprising because the CD44 receptor is known to be mechanosensitive. For example, a recent study showed that CD44 strengthens monoculture brain endothelial cell barrier function in the presence of intravascular fluid shear stress *in vitro*^[35]^. Moreover, multiple studies have reported that interstitial flow promotes cancer cell invasion via CD44-mediated mechanisms^[36]^. Yet, our results suggest that unlike in cancer cells, CD44 is not essentially involved in endothelial cell sprouting prompted by interstitial flow. In addition, we observed that broad spectrum inhibition of MMPs with GM6001 only partially inhibits interstitial flow-induced sprouting, thereby supporting a previous report that interstitial flow-induced sprouting is partially dependent on MMP activity^[8a]^. These results in presence of anti-CD44 and GM6001 inhibitor reinforce the potency of standalone interstitial flow in eliciting angiogenesis, particularly when flow is oriented against the direction of endothelial sprouting^[11-14]^.

In the absence of interstitial flow, we observe that ECM HA inhibits endothelial sprouting. This result is consistent with previous reports that under *in vitro* static conditions, addition of HA to collagen ECM inhibits capillary tube formation^[37]^ and enhances vascular barrier function^[35]^. Indeed, we previously reported that the addition of HA to collagen-based matrices suppresses both vessel sprouting and permeability under static *in vitro* conditions and in the absence of CXCL12 treatment^[21]^. One possible explanation for these observed decreases in angiogenic activity under static *in vitro* conditions is the addition of HA increases the solid fraction of collagen-based ECM hydrogels. Numerous *in vitro* studies have reported that under static conditions, increasing the hydrogel concentration of a single fibrillar ECM constituent (e.g., collagen or fibrin) results in decreased vessel outgrowth^[38]^. These outcomes have been attributed to steric hindrances due to increased density of matrix fibers that preclude ECM deformation, degradation, and remodeling that is necessary for vessel morphogenesis. It was also described that increasing fibrin ECM density limits vessel morphogenesis by hindering molecular diffusion from a proangiogenic source (stromal fibroblasts)^[38c]^. Yet, unlike with standalone collagen or fibrin ECM where there is an inverse relationship between ECM concentration and matrix pore size, we previously reported^[34]^, and confirmed in the present study, that addition of non-fibrillar HA increases fibrillar collagen matrix pore size due to swelling of this GAG constituent^[39]^. Collectively, these previous findings combined with the results from the present study suggest that the role of HA on angiogenesis is highly contingent on the presence or absence of a pro-angiogenic stimulus, whether biochemical or biophysical in nature.

To our knowledge, this report is the first to describe the important role of the ECM physical properties conferred by HA in regulating interstitial flow-mediated angiogenesis. We speculate that the significant increase in mean pore size of collagen/HA versus collagen-only matrices facilitates the initial extension and sustained elongation of HUVECs into the ECM when prompted to sprout by interstitial flow. It is important to recognize that while the mean pore size within the matrices used in this study are much smaller than the size of an individual cell, collagen fibers have been shown to be sufficiently deformable to create enough void spaces for endothelial sprout elongation^[40]^. It is also important to consider that our measurements for matrix microarchitecture were in acellular collagen-based hydrogels that reflect the initial ECM conditions prior to sprouting morphogenesis and matrix remodeling. Therefore, a limitation of our study is that we cannot account for the dynamic changes in matrix microarchitecture and stiffness that occur during endothelial sprouting and ECM remodeling. Moreover, we do not measure the complex and dynamic changes in flow induced shear stress, which depend strongly on the pore size between cells and matrix components and likely vary widely, even along individual cell membranes^[41]^. In the present study we observed more rapid and sustained interstitial flow induced endothelial sprouting in stiffer collagen/HA versus collagen-only matrices. Therefore, our results suggest that matrix pore size, and not matrix stiffness, is the primary ECM physical property that mediates interstitial flow induced endothelial sprouting. A recent study that used collagen-based microfluidic models to study interstitial flow mediated breast cancer cell invasion also concluded that matrix pore size, and not stiffness, was the primary determinant of cancer cell escape into a vascular-like cavity^[42]^. Thus, it would be interesting in future studies to investigate the combined effects of interstitial flow and ECM physical properties in orchestrating cancer cell-vascular interactions that lead to intravasation events and metastatic dissemination.

A key finding from our study is that HAdase treatment inhibits interstitial flow-promoted angiogenesis by altering the mechanical and microstructural properties of collagen/HA ECM. HAdase has been used in medical applications for decades^[43]^, primarily as an adjuvant to accelerate the absorption and dispersion of drugs into tissue, including HA-rich primary tumors^[44]^ and metastases^[45]^. In the context of neovascularization, HAdase is believed to be a pro-angiogenic stimulus by degrading angiostatic native high molecular weight HA into pro-angiogenic HA fragments^[46]^. However, the specific mechanisms driving these size-dependent effects of HA are largely unknown^[47]^. Our results suggest that HAdase may confer anti-angiogenic outcomes in HA-rich environments and in the presence of interstitial flow. Since tumor growth is angiogenesis-dependent^[48]^, immense research efforts have been directed towards developing anti-angiogenic therapies that directly neutralize the induction of angiogenesis by pro-angiogenic molecules, most prominently VEGF^[49]^. Yet, even when anti-angiogenic drugs have yielded significant clinical trial results leading to regulatory approval, improvements in patient survival have been disappointingly modest on the order of weeks to months^[50]^, primarily due to intrinsic/extrinsic resistance mechanisms^[51]^. The results from our study demonstrate biophysical induction of angiogenesis that is independent of biomolecular angiogenic stimulation and suggests that angiogenesis can be manipulated indirectly with enzymatic modifications to the ECM. Collectively, the findings from our study points to the need for further investigation in the role of ECM physical properties in regulating angiogenesis and anti-angiogenesis mechanisms.

## 4. Conclusion

Using 3D microfluidic biomimicry with detailed characterization of hydrogel scaffolds, we investigated how ECM physical properties influence interstitial flow-mediated sprouting angiogenesis. We observed that the addition of HA to collagen-based hydrogels significantly enhances the initial extension and sustained elongation of endothelial sprouting in response to interstitial flow. Interestingly, we show that enhancement in interstitial flow-promoted sprouting in collagen/HA matrices was inhibited significantly with physical remodeling of ECM HA by HAdase but not with blocking HA-CD44 interactions. We also show that HA significantly increases ECM pore size and stiffness of collagen-based matrices. Collectively, these novel findings comprise an important advancement towards refining our understanding of angiogenesis that is mediated by the biophysical microenvironment. Moreover, these results will help inform future developments in pro-angiogenic biomaterials and *in vitro* disease models (e.g., organs-on-a-chip) for interrogating mechanisms of vessel outgrowth and remodeling.

## 5. Materials and Methods

### 5.1 Fabrication of microfluidic device

The microfluidic microvessel analogue was fabricated using PDMS lithography. Briefly, the basic and curing agent of PDMS precursor was mixed at 10:1 ratio and poured onto a patterned mold wafer. Then, the patterned PDMS layer was irreversibly bonded onto a glass slide by plasma oxidation treatment. The assembled microfluidic device was then placed in the 65 °C overnight to promote the bonding between PDMS and glass slide. The dimensions of channels are 500, 300 and 100 μm in width for the vascular channel, ECM compartment and fluidic channel, correspondingly. The height of engineered microvessel analogue is ∼ 70 μm. Type I collagen gel (Corning Inc.) isolated from rat tail was introduced into the central ECM compartment and polymerized at 37 °C in a humidified incubator overnight prior to cell seeding. HA (purchased from Sigma) was mixed with collagen gel at a concentration of 1 mg/mL HA to prepare collagen/HA matrices. The collagen-only or the collagen/HA pre-polymer mixture was pipetted into the central channel and polymerized overnight in a humidified incubator before further experimental usage.

### 5.2 Biophysical characterization of extracellular cellular matrices

The mechanical stiffness of ECM hydrogels was measured using the protocols previously reported by our group^[21, 34]^. Briefly, 300 µL of 3 mg/mL concentration of collagen-only and collagen/HA were casted on the 10.4 mm diameter custom made polystyrene housing holder. Biochemical treatments including inhibition of MMP activities (20 µM GM-6001) and enzymatic matrix degradation of HA (1mg/mL hyaluronidase) was performed by incubating the ECM hydrogels with 200 μl of individual reagents for one day in a humidified incubator. Samples were kept immersed in phosphate-buffered saline (PBS) during mechanical testing to maintain the collagen-based gels in a hydrated environment. Stiffness measurement was conducted by the high precision indentation system which was programmed to indent the gel to up to 40% strain at 10 % increasing intervals lasting 300 seconds each. The peak loads responses were automatically recorded and used to calculate stress using the displacement at each interval and known indenter geometry (diameter of 4.8 mm). Stress responses were automatically recorded corresponding to each strain. The indentation modulus is defined as the ratio of stress over strain.

### 5.3 Collagen fiber imaging and mean pore size analysis

The collagen fiber of both collagen-only gel and collagen/HA in the microvessel analogues were imaged by imaged using confocal microscopy on a Nikon A1R Live Cell Imaging Confocal Microscope via a 40X 1.3 NA oil immersion lens controlled with NIS-Elements software. Confocal reflectance stacks of approximately 50 µm in height with a z-step of 0.59 µm were acquired for image analysis (80-90 total images per stack). When imaging, 3 stacks per devices were obtained. The average pore size of the matrix was estimated by use of the MO Method^[52]^. In this approach, the average pore size was quantified by morphological operations on binary images was used using a house-made MATLAB script.

### 5.4 Characterization of interstitial flow

To generate interstitial flow, pipettes tips with 80 µL culture media were inserted to the two inlets of fluidic channel. Texas-red conjugated Dextran fluorescent dye was introduced in the same manner as the interstitial flow setup in the cell culture experiments. Time-lapse fluorescent microscope was employed to track the flow of dye into the collagen gel over time. Then, we applied the control volume approach of fluid flux transport to evaluate the average flow velocity. The simplified advection transport equation is

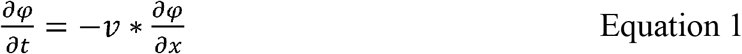

where ψ, v, t, x are the fluorescent intensity of dye, average flow velocity, time, and distance along flow direction, respectively. The assumptions here are: (i) flow in the microfluidic device is one-dimensional (i.e., flow across the ECM compartment is only present in the direction perpendicular to channels), and (ii) convective transport is dominant in interstitial flow condition. Using this method, interstitial flow velocity in the microdevice with the vascular channel lined with HUVECs was ∼ 10.76 and 11.18 µm/s for microvessel of collagen-only and collagen/HA matrix, correspondingly.

### 5.5 Preparation of HUVECs

Human umbilical vein endothelial cells (HUVECs) were purchased from Lonza and cultured in endothelial growth medium 2 (EGM-2) medium (Lonza). HUVECs were cultured in T-75 flasks in a humidified cell culture incubator at 37 °C and 5 % CO_2_ with culture medium being changed every two days. Cell passage numbers of 5 - 12 were used in this study. To improve the biocompatibility of the microfluidic device, the vascular channels were coated with fibronectin solution at a concentration of 100 μg/mL at least 30 minutes prior to cell seeding. HUVECs were harvested from a flask using 0.05 % EDTA-Trypsin (Invitrogen) for 4 minutes, and the cell suspension was centrifuged at 950 RPM for 4 minutes. To acquire a monolayer of HUVECs lining the vascular channels, the cell suspension was adjusted to ∼ 2 × 10^7^ cells/mL and introduced to the channel by pipetting. Medium was changed every day in the device to ensure better cell attachment and healthy cell growth. For all experiments, treated conditions were introduced after cells had been cultured in the microfluidic device for 1 day to ensure treatment had no effect on initial cell attachment. For CD44 blocking experiments, trypsinized HUVECs were treated with medium containing CD44 antibody (2 μg/mL, MA5-13890, Invitrogen) for 30 minutes at room temperature prior to seeding. GM6001 (CC10, Millipore Sigma) was diluted to 20 µM in culture medium and introduced to HUVECs cultured in microvessel analogue for the MMP inhibition experiments. For HAdase treatment, HAdase was diluted to 1mg/mL in the culture medium and applied to HUVECs cultured in microvessel analogue. To generate the interstitial flow, pipet tips with 80 µl medium were inserted to the two inlets of fluidic channels. The inserted pipet tips were replenished every day to ensure the pressure head was maintained.

### 5.6 Sprouting angiogenesis assay and sprouting elongation analysis

HUVECs were cultured in microfluidic devices for three days using endothelial cell culture medium. Phase-contrast images of sprouts from ECM/vessel interfaces were taken every 24 hours and a house-made MATLAB code was used to evaluate the specific sprouting area of each device. To evaluate the elongation of angiogenic sprouting, ECM compartment was divided into 10 equal regions and a house-made MATLAB code was applied to analyze total sprouting area in each ECM regions to generate a distribution of angiogenic sprouting area across entire ECM compartment. To analyze sprouting number in MMP inhibition, the normalized sprouting number percentage is defined as a percentage of the total sprouting number per device divided by total number of HUVECs/ECM interface in a device.

### 5.8 Immunofluorescence staining

HUVECs cultured in the microvessel analogues were fixed by 4% paraformaldehyde (Sigma-Aldrich, 158127) for 15 min at room temperature. Then, the microvessels were permeabilized by incubating with 0.1% Triton X-100 (Sigma-Aldrich, T9284) for 10 min. Subsequently, the microvessels were incubated with 0.1% bovine serum albumin (Sigma, A7906) overnight at 4 °C to avoid nonspecific binding. VE-cadherin was stained by incubation with VE-Cadherin primary antibody (Santa Cruz Biotechnology, SC-9989) at 1:125 dilution in DPBS overnight and washed with 1% Tween-20 (Bio-Rad Laboratories, 1610781) subsequently. HUVECs were then stained with Alexa Fluor 488 donkey second antibody (Invitrogen, A21202) at 1:500 dilution in DPBS for 3 hours at room temperature. F-actin is labeled with Alexa Fluor 546 Phalloidin (Invitrogen, A22283) at 1:100 in DPBS for 3 hours at room temperature. Cell nuclei of HUVECs were stained by 4′,6-diamidino-2-phenylindole (DAPI) (Invitrogen, D1306) at 1:500 dilution for 15 min. Confocal microscopy was performed on the stained microvessel using a broadband confocal microscope (TCS SP5, Leica Microsystems, Wetzlar, Germany).

### 5.7 Statistical analysis

Numerical data reported in this manuscript were expressed as mean ± the standard error of mean (S.E.M). Each experimental condition was performed at least in three replicates to conduct statistical analysis. Variations of all data were statistically analyzed by performing one-way ANOVA followed by post-hoc unpaired, two-tailed Student t test, executed by origin lab software. To compare the statistical difference of each experimental condition, the asterisk mark (*) was applied as * for p-value <0.05, ** for p-value <0.01, and *** for p-value <0.001.

## Conflicts of Interest

The authors declare that they have no competing interests.

## Acknowledgements

This work was supported by funding awarded to J.W.S. from an NSF CAREER Award (CBET-1752106), The American Heart Association (15SDG25480000), Pelotonia Junior Investigator Award, NHLBI (R01HL141941), and The Ohio State University Materials Research Seed Grant Program, funded by the Center for Emergent Materials, an NSF-MRSEC, grant DMR-1420451, the Center for Exploration of Novel Complex Materials, and the Institute for Materials Research. This work was also supported by funding awarded to Y.-C.T. from Taiwan Ministry of Science and Technology (MOST 109–2221-E-001–002-MY2 and 110–2221-E-001–005-MY3). Partial personnel support through The Mark Foundation for Cancer Research (18-024-ASP) is also acknowledged. C.-W.C, P.E.B, and A.A. gratefully acknowledge funding from the Pelotonia Graduate Fellowship Program. M.C.-M. thanks the supports from an OSU Graduate Enrichment Fellowship, a Discovery Scholars Fellowship, and a NHLBI Diversity Supplement.

## Notes

### Competing Interest Statement

The authors have declared no competing interest.

