## Supplemental Information for "ECM-derived biophysical cues mediate interstitial flow-induced sprouting angiogenesis"

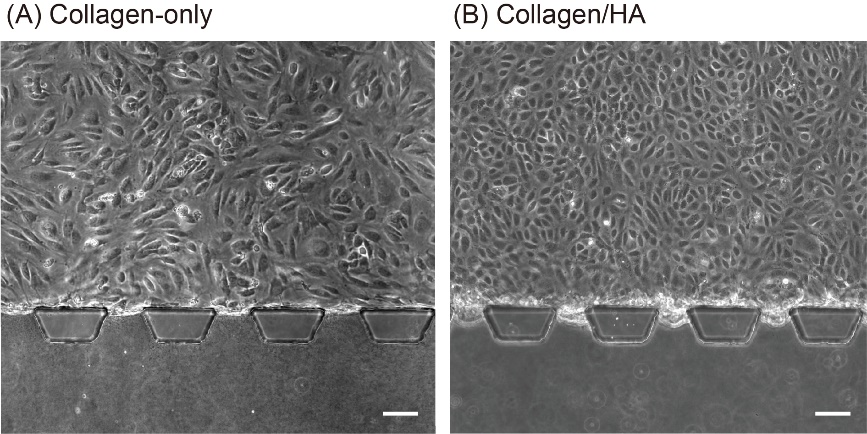

**Figure S1.** Endothelium of HUVEC in the microfluidic microvessel analogue under static condition on day 3. The phase contrast image of microvessels in (A) collagen-only and (B) collagen/HA ECM. (Scale bars = 50 µm)

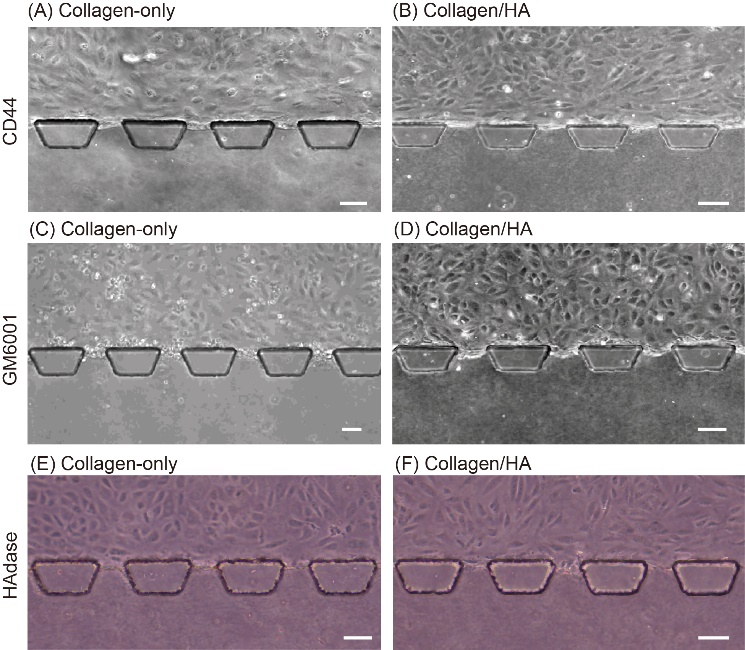

**Figure S2.** Endothelium of HUVECs in the microfluidic microvessel analogue in responses of anti-CD44 (A and B), GM6001 (C and D), and HAdase (E and F) treatments under static condition on day 1. (Scale bars = 50 µm)

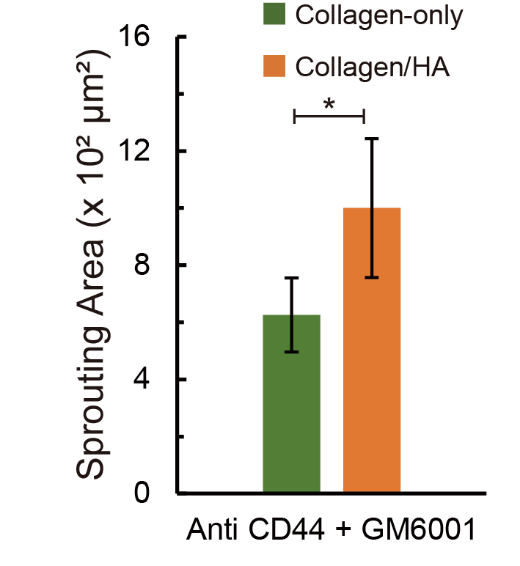

**Figure S3.** Sprouting area of HUVECs cultured in microvessel analogues on day 1 in response to combined treatment of CD44 blocking and GM6001 under interstitial flow. The data were expressed as mean ± standard error of mean (n ≥ 3). One-way ANOVA followed by post-hoc unpaired, two-tailed Student t test was performed to evaluate the statistical significance. * indicates p-value < 0.05.
